# Glucosyltransferase-dependent and independent effects of *Clostridioides difficile* toxins during infection

**DOI:** 10.1101/2021.09.22.460915

**Authors:** F. Christopher Peritore-Galve, John A. Shupe, Rory J. Cave, M. Kay Washington, Sarah A. Kuehne, D. Borden Lacy

**Affiliations:** Department of Pathology, Microbiology, and Immunology, Vanderbilt University Medical Center, Nashville, Tennessee, USA, 37232; School of Biomedical Sciences, University of West London, London, United Kingdom, TW8 9GB; Oral Microbiology Group, School of Dentistry and Institute of Microbiology and Infection, College of Medical and Dental Sciences, The University of Birmingham, Birmingham, United Kingdom, B5 7EG; Department of Veterans Affairs Medical Center, Nashville, Tennessee, USA, 37212

## Abstract

*Clostridioides difficile* infection (CDI) is the leading cause of nosocomial diarrhea and pseudomembranous colitis in the USA. In addition to these symptoms, patients with CDI can develop severe inflammation and tissue damage, resulting in life-threatening toxic megacolon. CDI is mediated by two large homologous protein toxins, TcdA and TcdB, that bind and hijack receptors to enter host cells where they use glucosyltransferase (GT) enzymes to inactivate Rho family GTPases. GT-dependent intoxication elicits cytopathic changes, cytokine production, and apoptosis. At higher concentrations TcdB induces GT-independent necrosis in cells and tissue by stimulating production of reactive oxygen species via recruitment of the NADPH oxidase complex. Although GT-independent necrosis has been observed *in vitro*, the relevance of this mechanism during CDI has remained an outstanding question in the field. In this study we generated novel *C. difficile* toxin mutants in the hypervirulent BI/NAP1/PCR-ribotype 027 R20291 strain to test the hypothesis that GT-independent epithelial damage occurs during CDI. Using the mouse model of CDI, we observed that epithelial damage occurs through a GT-independent process that is does not involve immune cell influx. The GT-activity of either toxin was sufficient to cause severe edema and inflammation, yet GT activity of both toxins was necessary to produce severe watery diarrhea. These results indicate that both TcdA and TcdB contribute to infection when present. Further, while inactivating GT activity of *C. difficile* toxins may suppress diarrhea and deleterious GT-dependent immune responses, the potential of severe GT-independent epithelial damage merits consideration when developing toxin-based therapeutics against CDI.

**SIGNIFICANCE:** *Clostridioides difficile* is the leading cause of antibiotic-associated diarrhea in hospitals worldwide. This bacterium produces two virulence factors, TcdA and TcdB, which are large protein toxins that enter host colon cells to cause inflammation, fluid secretion, and cell death. The enzymatic domain of TcdB is a target for novel *C. difficile* infection (CDI) therapeutics since it is considered the major factor in causing severe CDI. However, necrotic cell death due to non-enzymatic TcdB-host interactions have been reported in cell culture and intoxicated tissue. Here, we generated *C. difficile* strains with enzyme-inactive toxins to evaluate the role of each toxin in an animal model of CDI. We observe an additive role for TcdA and TcdB in disease and both glucosyltransferase-dependent and independent phenotypes. These findings are expected to inform the development of toxin-based CDI therapeutics.

## INTRODUCTION

*Clostridioides difficile* infection (CDI; formerly *Clostridium difficile*) is the leading cause of hospital-acquired diarrhea and pseudomembranous colitis in the USA (1, 2). *C. difficile* is a Gram-positive, spore-forming anaerobe that infects the colon, causing mild to severe symptoms including diarrhea, pseudomembranous colitis, toxic megacolon, and in severe cases, death (3). CDI is prevalent among elderly and immunocompromised individuals in healthcare settings, typically following treatment with broad spectrum antibiotics. However, the rate of community-acquired infections among healthy individuals has increased over the past two decades due to the emergence of novel epidemic *C. difficile* strains (3, 4). Despite the clinical importance of CDI, we do not have a complete understanding of molecular host-microbe interactions during infection, which hampers progress towards developing effective prevention and treatment strategies.

CDI is mediated by two large homologous protein toxins, TcdA (308 kDa) and TcdB (270 kDa) that are secreted during infection to cause disease symptoms. The toxins bind host cell receptors and become internalized into vesicles via endocytosis (5). Endosome acidification elicits structural changes in the toxins, stimulating pore formation and translocation of the N-terminal glucosyltransferase (GT) and autoprocessing (AP) domains into the cytosol (5). Inositol-6-phosphate-induced autoprocessing releases the GT domain in the host cell and permits access to Rho family GTPases. GT activity transfers glucose from UDP-glucose onto Rho GTPases, irreversibly inactivating these regulatory proteins (5–8). This inactivation causes cytoskeletal rearrangement, leading to the disruption of cell adhesion junctions, cytopathic changes, and apoptosis (5). These effects stimulate proinflammatory cytokines and neutrophil chemoattractants that generate acute inflammatory responses. Prolonged host inflammation during CDI increases the severity of tissue damage and the probability of lethal disease outcomes (9–11).

In addition to GT-dependent effects of *C. difficile* toxins, TcdB can induce GT-independent necrotic cell death at high concentrations *in vitro* (> 0.1 nM) (12, 13). This effect occurs through the stimulation of reactive oxygen species via the NADPH oxidase complex (14). In contrast, TcdA induces GT-dependent cell death through the apoptosis pathway at both high and low concentrations (15). TcdA and TcdB can also stimulate cytokine release *in vitro* through both GT-dependent and independent pathways (9). The relevance of GT-independent cell death during infection has been unclear since it is unknown if cells encounter the concentration of TcdB required to stimulate necrotic cell death. A recent study concluded that GT activity was necessary to cause CDI in mouse and hamster models of infection but does not preclude an additional role for GT-independent events as the infection progresses (16). The study also involved GT-defective *C. difficile* mutants in the 630 strain, which is genetically tractable, but causes few disease symptoms compared to BI/NAP1/PCR-ribotype 027 epidemic strains (16, 17).

Historically, TcdA was considered the toxin responsible for CDI pathogenesis (18). The identification of patient-derived clinical isolates that are TcdA-negative and TcdB-positive, along with functional analyses of toxin mutants in animal models indicated however that TcdB is necessary and sufficient to cause severe CDI (19, 20). Other data demonstrate that TcdA alone can cause symptoms in the hamster model of infection, suggesting a potential role for both toxins during disease (21). Overall, the respective roles of TcdA and TcdB during infection are not fully understood.

CDI remains a pervasive disease across the world and poses a significant burden on patients and the healthcare system. The main goal of this study was to define GT-dependent and independent effects of each toxin during CDI using the mouse model of infection and a BI/NAP1/PCR-ribotype 027 epidemic strain. The results from this study provide a framework to further understand the complex molecular interplay between *C. difficile* toxins and the host colon during infection.

## RESULTS

### Generation of glucosyltransferase-deficient mutants and *in vitro* characterization

To define GT-dependent and independent effects of TcdA and TcdB during infection, GT-deficient mutants were generated in the epidemic *C. difficile* BI/NAP1/PCR-ribotype 027 R20291 background (hereafter referred to as R20291) using homologous allelic exchange (22). This approach allowed for the introduction of SNMs in the GTD of either or both toxins, causing single point mutations at positions TcdA::D285N/D287N and/or TcdB::D286N/D288N (Fig 1A). Four novel mutant strains were created, including GT-deficient mutants for each toxin (A_GTX_ B+ and A+ B_GTX_), a GT-deficient mutant of both toxins (A_GTX_ B_GTX_), and a total *tcdA* knockout with GT-deficient TcdB (Δ*tcdA* B_GTX_) (Table 1). Whole genome sequencing was performed on all strains used in this study to ensure the proper mutations were present and that there were no off-target mutations that might affect their phenotypes (Table 1; Fig 1A) (23). There were no unexpected variations in nucleotides for any of the mutant strains compared to the wildtype background and the reference R20291 genome (NC_013316). All sequencing reads are available in the NCBI Sequencing Read Archive under accession number PRJNA762329.

**TABLE 1.** Strains and plasmids used in this study.

| Strains | Description | Source or Citation |
| --- | --- | --- |
| *Clostridioides difficile* |  |  |
| R20291Δ*pyrE* | Wildtype BI/NAP1/PCR-ribotype 027 strain obtained from CRG0825 stock at the Nottingham Clostridia Research Group, Cardiff, UK with *pyrE* knocked out. | (22) |
| R20291 | Wildtype BI/NAP1/PCR-ribotype 027 strain obtained from CRG0825 stock at the Nottingham Clostridia Research Group, Cardiff, UK with restored *pyrE*. | (22) |
| A<sub>GTX</sub> B+ | R20291 TcdA::D285N/D287N | This study |
| A+ B<sub>GTX</sub> | R20291 TcdB::D286N/D288N | This study |
| A<sub>GTX</sub> B<sub>GTX</sub> | R20291 TcdA::D285N/D287N-TcdB::D286N/D288N | This study |
| Δ*tcdA* B<sub>GTX</sub> | R20291 Δ*tcdA* TcdB::D286N/D288N | This study |
| Δ*tcdA* Δ*tcdB* | R20291 *tcdA/tcdB* ClosTron insertional mutants | (44) |
| *Escherichia coli* |  |  |
| DH5α | Chemically competent *E. coli* cells for transformation | NEB |
| CA434 | Conjugation host | (35) |
| Plasmids | Description | Source or Citation |
| pMTL-YN4 | Knockout vector for R20291 | (22) |
| pMTL-YN4::TcdA | KO vector backbone with 2393 bp fragment of *tcdA* GTD | This study |
| pMTL-YN4::TcdB | KO vector backbone with 2332 bp fragment of *tcdB* GTD | This study |
| pMTL-YN4::TcdA D285N/D287N | KO vector backbone with mutations in *tcdA* | This study |
| pMTL-YN4::TcdB D286N/D288N | KO vector backbone with mutations in *tcdA* | This study |
| pMTL-YN4::TcdA KO | KO vector backbone with left flank and right flanks of *tcdA* for full gene deletion | This study |

**FIGURE 1.**
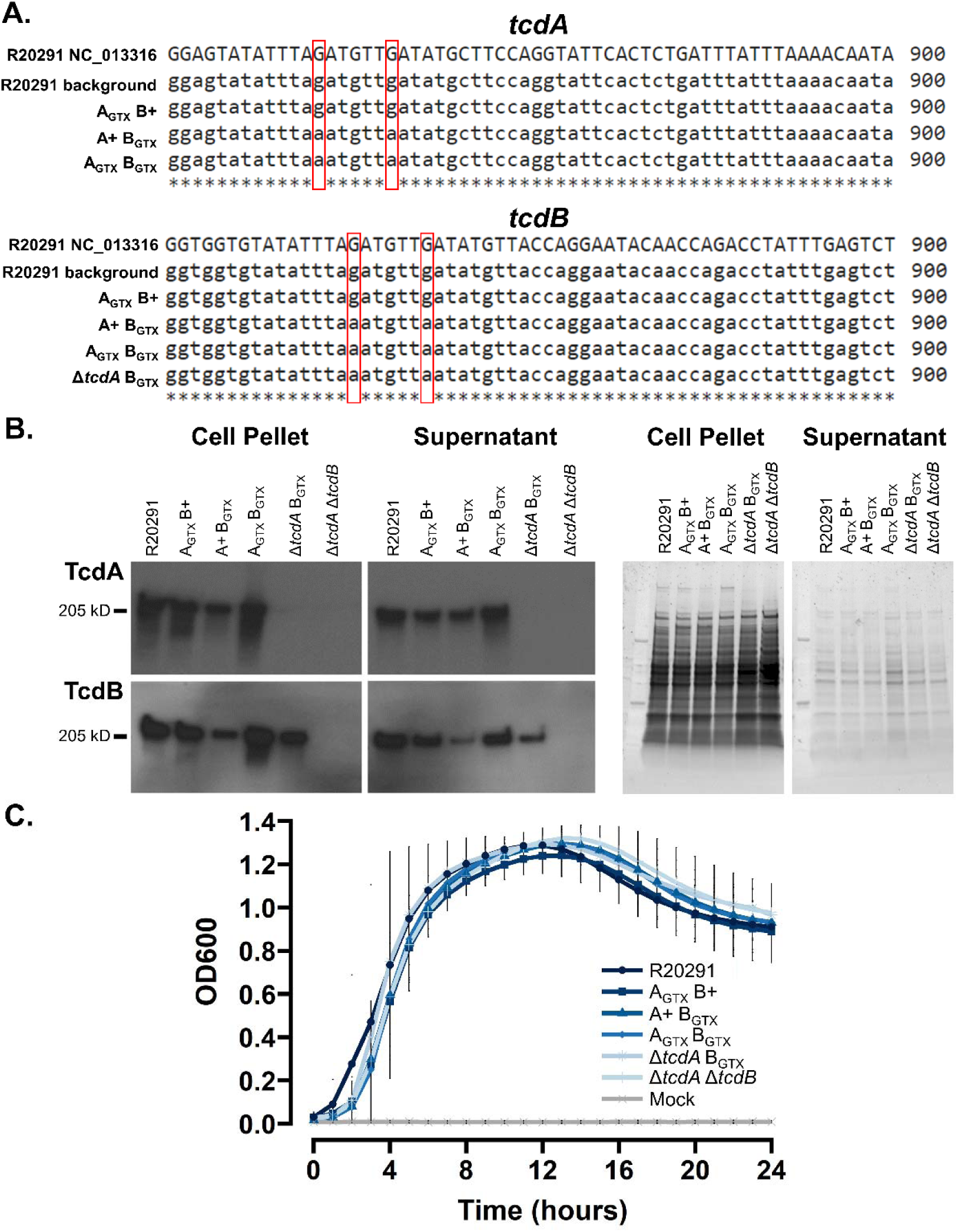
*In vitro* characterization of *C. difficile* mutants. **(A) *tcdA* and *tcdB* gene alignments of mutant and wildtype strains to the reference genome of R20291**. Red boxes highlight SNMs that inactivate GT catalytic activity. (B) Western blot images of mutant and wildtype strains to confirm production and secretion (or lack thereof) of TcdA and TcdB. Images on right are SDS-PAGE gels stained with SYPRO ruby to visualize total protein loaded for Western blots. (C) *In vitro* growth curves of each strain and mock-inoculated controls (*n* = 5 per treatment) cultured in BHIS for 24 hours. Dots are the average at each given timepoint, and the error bars depict the 95% confidence interval.

The novel strains were tested for toxin production and bacterial growth *in vitro* to ensure that the introduced mutations had no unintended effects on these processes. Toxin production and secretion was determined in mutant and wildtype strains cultured for 24 hours in toxin-inducing TY medium. After 24 hours of growth, cell pellets and supernatants were separated by centrifugation, then supernatants were filter-sterilized to remove any cell contaminants. Immunoblot analyses of TcdA and TcdB determined that there were no GT-dependent effects on toxin production or secretion and confirmed the knockout phenotypes of Δ*tcdA* B_GTX_ and Δ*tcdA* Δ*tcdB* (Fig 1B).

*In vitro* growth of each strain was assessed in BHIS medium for 24 hours under anaerobic conditions. Bacterial growth was measured every hour and revealed that there were no significant differences in growth between mutant and wildtype strains (Fig 1C). Validation of mutant genotypes, toxin production and secretion, and bacterial growth was essential to proceeding with animal infection experiments and to demonstrate that *in vivo* phenotypes are not due to detectable physiological defects in mutant strains.

### Glucosyltransferase activity of both toxins causes the most severe disease outcomes

The non-lethal mouse model of infection was used to define GT-dependent and independent effects of each toxin during CDI. Mice were pre-treated with cefoperazone antibiotics in drinking water for five days, then were returned to regular drinking water for two days prior to inoculation with *C. difficile* (Fig 2A). *C. difficile* spores of each strain (10^5^ CFU/mL) were administered via oral gavage, and metrics of infection and symptom severity were assessed daily until 2- or 4-days post-inoculation (dpi). The most severe decline in animal weight occurred at 2 dpi, then mice began to recover by 4 dpi (Fig 2B). Thus, the starkest differences between treatments were observed at 2 dpi (Sup Fig 1). At 2 dpi, the wildtype R20291 strain induced 12% weight loss on average, which was significantly more than all other groups, even when compared to A_GTX_ B+ (*p* = 0.0075) and A+ B_GTX_ (*p* < 0.0001) (Fig 2B and Sup Fig 1). A_GTX_ B+ caused the second highest average weight loss at 2 dpi (8.9%), but was not significantly different from A+ B_GTX_, which caused 6.4% average weight loss (*p* = 0.2273; Fig 2B and Sup Fig 1). Mice inoculated with A_GTX_ B_GTX_, Δ*tcdA* B_GTX_, Δ*tcdA* Δ*tcdB*, or mock (sterile PBS) lost little to no weight (Fig 2B and Sup Fig 1). Similar trends between treatments were observed at 3 and 4 dpi (Fig 2B and Sup Fig 1).

**FIGURE 2.**
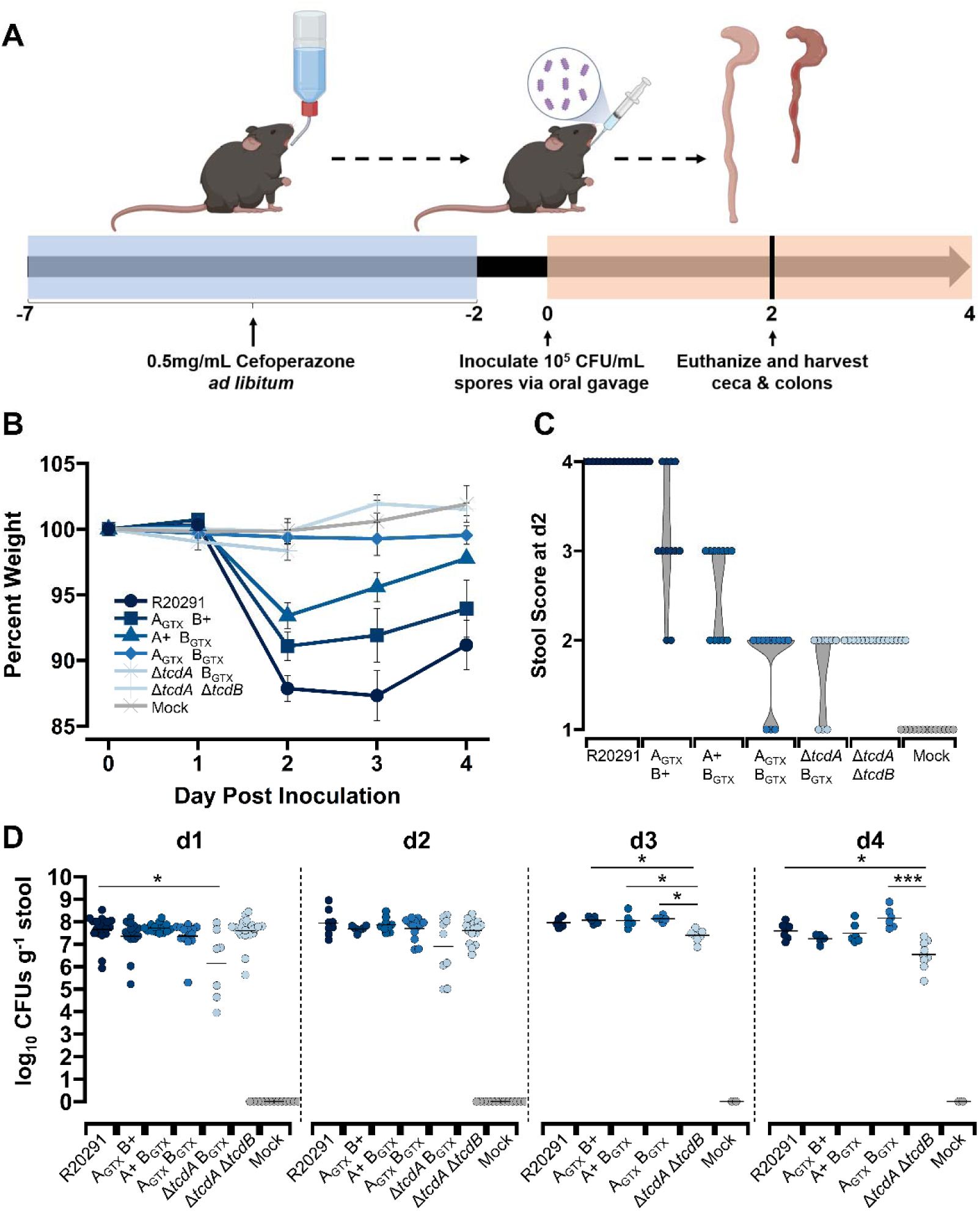
GT-dependent and independent effects on weight loss, diarrhea, and colonization in the mouse model of *C. difficile* infection. (A) Visual abstract of the cefoperazone mouse model of CDI used for this study. (B) Percent weight loss from day 0 for R20291 (*n* = 22), A_GTX_ B+ (*n* = 15), A+ B_GTX_ (*n* = 15), A_GTX_ B_GTX_ (*n* = 15), Δ*tcdA* B_GTX_ (*n* = 9), Δ*tcdA* Δ*tcdB* (*n* = 22), and mock (*n* = 13). Points represent group averages at each day and error bars denote standard error of the mean. (C) Stool scores at 2 dpi. (D) Daily *C. difficile* burden in stool. Each point is an individual mouse, and the crossbars represent group daily means. Significantly different groups as determined by Dunn’s test are shown in brackets (* *p* < 0.05; *** *p* < 0.001).

Weight loss is typically caused by increased fluid loss during CDI-associated diarrhea. To visually test the effects of GT activity of each toxin on diarrhea severity, mouse stool was collected daily for visual assessment of moisture, color, and consistency. Stool scores from 1-to-4 were assigned based on pre-defined criteria. All mice inoculated with wildtype R20291 had a stool score of 4 at 2 dpi based on the presence of watery diarrhea and wet tail (Fig 2C). Although some mice inoculated with A_GTX_/B+ developed severe watery diarrhea, most had a stool score of 3, characterized by soft and discolored stool (Fig 2C). No mice inoculated with A+/B_GTX_ developed watery diarrhea, and typically displayed soft, discolored stool (Fig 2C). Stool from mutant strains that induced no weight loss (A_GTX_ B_GTX_, Δ*tcdA* B_GTX_, and Δ*tcdA* Δ*tcdB*) was well formed, yet discolored (Fig 2C).

To quantify *C. difficile* burden during infection, daily fecal samples were collected, weighed, homogenized, then dilution plated onto semi-selective media. At 1 dpi, strains colonized to a high density (∼10^7^ CFU g^-1^ stool), however, R20291 was significantly more abundant in stool compared to Δ*tcdA* B_GTX_ (*p* = 0.031; Fig 2D). There were no significant differences in *C. difficile* stool burden between strains at 2 dpi, when symptoms were most severe. Colonization density of Δ*tcdA* Δ*tcdB* began to significantly reduce at 3 and 4 dpi, while titers of other strains remained more consistent. As expected, no mock-inoculated mice contained *C. difficile* (Fig 2D). Since there were no notable differences in *C. difficile* burden in stool, it is expected that symptom severity phenotypes caused by each strain are due to molecular toxin-host interactions and not colonization burden.

### Edema and inflammation are glucosyltransferase-dependent, but epithelial damage is glucosyltransferase-independent

To visualize diarrhea and inflammation on a macroscale, mice were euthanized at 2 dpi, then ceca and colons were excised and imaged (*n* = 6 per treatment). The ceca and colons of mice inoculated with the wildtype R20291 strain were severely inflamed and contained very little wet stool (Fig 3A). Soft, discolored stool was typically observed in ceca and colons from mice inoculated with A_GTX_ B+ and A+ B_GTX_ (Fig 3A). Stool discoloration from A_GTX_ B_GTX_, Δ*tcdA* B_GTX_, and Δ*tcdA* Δ*tcdB*-inoculated mice was noted in ceca and colons, relative to well-formed, normal colored stool from mock-inoculated mice (Fig 3A).

**FIGURE 3.**
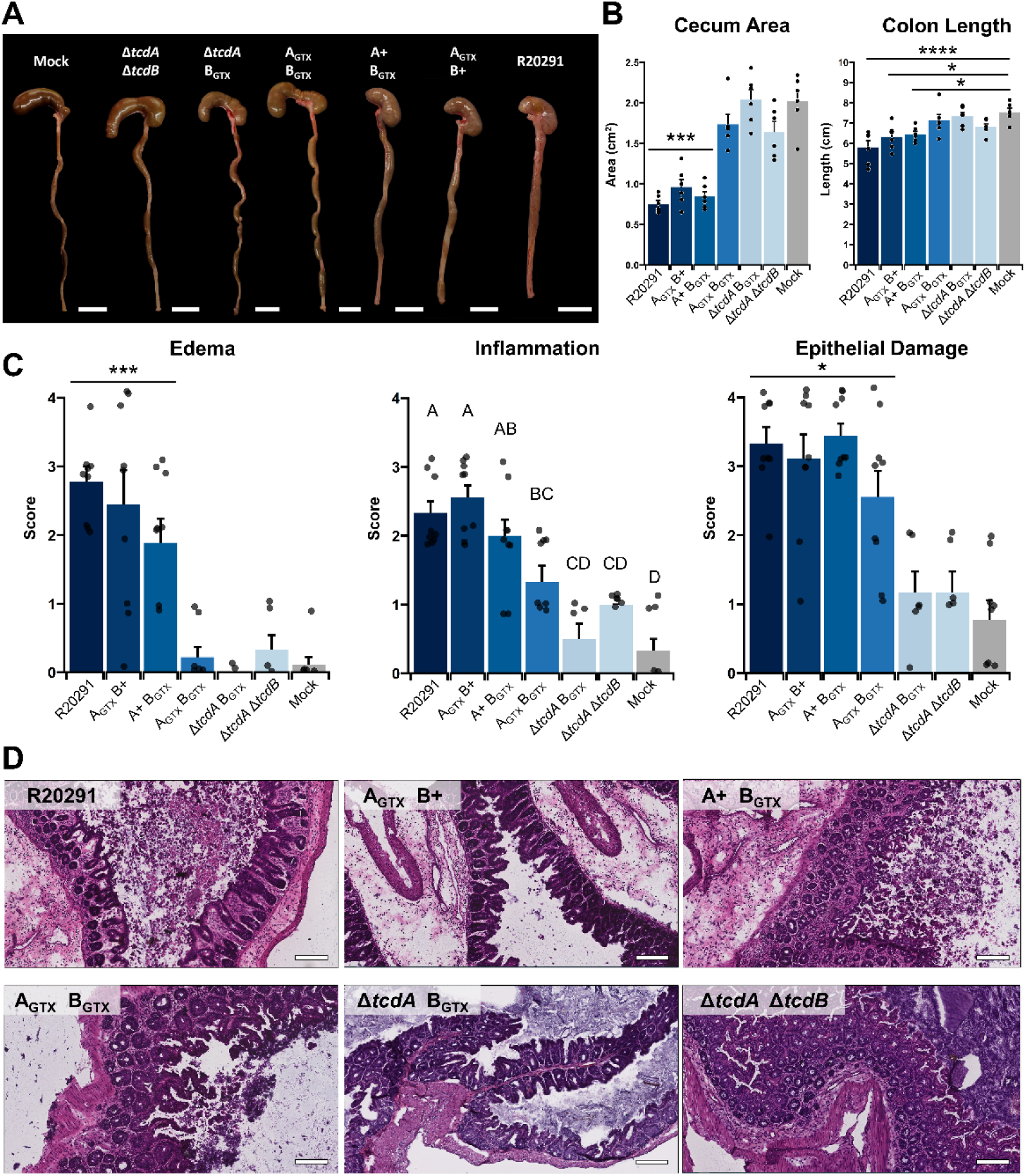
Glucosyltransferase activity causes edema and inflammation, but epithelial damage is glucosyltransferase independent. (A) Representative images of ceca and colons from mice inoculated with each treatment at 2 dpi. White bars denote 1 cm. (B) Cecum area and colon length as metrics of organ inflammation. Bars are the mean of each group and points are individuals within the group. Error bars represent standard error of the mean. Differences in cecum area depict that R20291, A_GTX_ B+, and A+ B_GTX_ are significantly smaller than the four other groups. Differences in colon length are comparing R20291, A_GTX_ B+, and A+ B_GTX_ to mock-inoculated colons. (C) Epithelial damage, inflammation, and edema mean scores as determined by a gastrointestinal pathologist. Points represent each individual animal and error bars are the standard error of the mean (*n* = 6-9 per treatment). Statistical differences as determined by Tukey’s HSD are shown in brackets (* *p* < 0.05; *** *p* < 0.001; **** *p* < 0.0001) or by letters (*p* < 0.05). (D) Representative H&E images of mouse ceca inoculated with each strain at 2 dpi. Scale bar, 80 μm.

Whole organ inflammation was determined by measuring cecum area and colon length (Fig 3B). The area of ceca from mice inoculated with R20291, A_GTX_ B+, and A+ B_GTX_ were half the size of those from mice inoculated with A_GTX_ B_GTX_, Δ*tcdA* B_GTX_, Δ*tcdA* Δ*tcdB*, and the mock control (*p* < 0.001; Fig 3B). Colon length was altered in response to weight loss-inducing strains, albeit less drastically than differences observed in ceca (Fig 3B). Colons from mice inoculated with R20291, A_GTX_ B+, and A+ B_GTX_ were significantly shorter than mock-inoculated mice (*p* < 0.05; Fig 3B). However, none were significantly shorter than mice inoculated with Δ*tcdA* Δ*tcdB* (*p* > 0.05; Fig 3B).

To examine histopathological phenotypes in infected tissues, excised ceca were fixed and frozen in OCT media, then cryosectioned and H&E stained. Edema, inflammation, and epithelial damage were scored on a 1-4 scale by a board-certified gastrointestinal pathologist based on previously published criteria (17). Cecal edema was observed in mice inoculated with R20291, A_GTX_ B+, and A+ B_GTX_, and was scored on average between 2 and 3 based on the presence of moderate to severe edema with widespread multifocal submucosal expansion (*p <* 0.001; Fig 3C and 3D) (17).

Phenotypes of cecal inflammation were more nuanced than edema. Once again, R20291, A_GTX_ B+, and A+ B_GTX_, caused the highest average inflammation scores (2-3), hallmarked by moderate to severe neutrophilic inflammation and submucosal to mural involvement (*p <* 0.001; Fig 3C and 3D) (17). However, A_GTX_ B_GTX_ caused minimal to moderate inflammation and did not significantly differ from inflammation induced by A+ B_GTX_ (*p* = 0.1736; Fig 3C and 3D).

Finally, epithelial damage was assessed. Severe epithelial damage was observed in mice inoculated with R20291, A_GTX_ B+, A+ B_GTX_, and A_GTX_ B_GTX_ (Fig 3C and 3D). Mice with epithelial damage scores between 3-4 exhibited severe multifocal epithelial vacuolation, apoptotic figures, and severely damaged regions developed pseudomembranes (Fig 3D and Fig 4) (17). Minimal epithelial damage was observed in Δ*tcdA* B_GTX_, Δ*tcdA* Δ*tcdB*, or mock-inoculated mice (Fig 3D and Fig 4).

**FIGURE 4.**
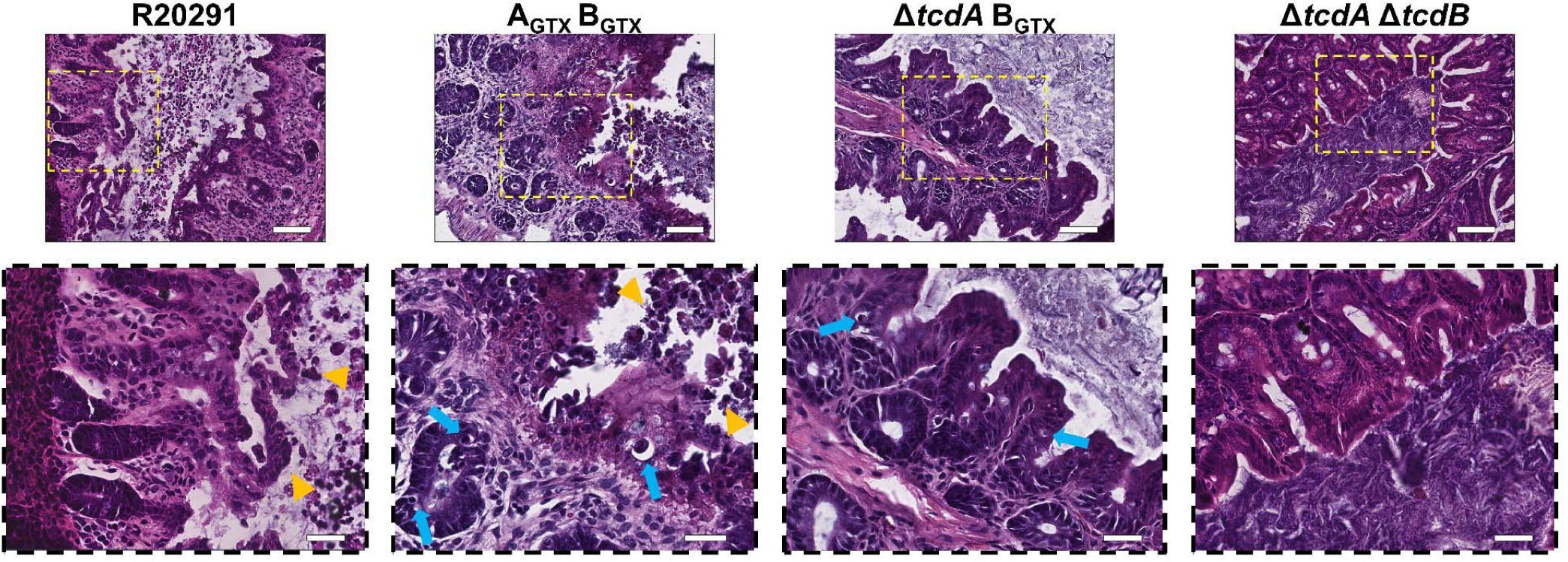
Epithelial damage is elicited by a glucosyltransferase-independent mechanism during infection. Representative images of severe epithelial injury in R20291 and A_GTX_ B_GTX_ compared to Δ*tcdA* B_GTX_ and Δ*tcdA* Δ*tcdB*. Scale bar, 80 μm in 20x magnification images. Zoomed in (40x magnification; dashed boxes) images offer a closer view at specific epithelial damage phenotypes. Apoptotic bodies or dying cells are highlighted by blue arrows. Sloughing dead cells and pre-pseudomembrane formation is shown by orange arrowheads. Scale bar, 130 μm.

### Glucosyltransferase activity of TcdB is required to elicit MPO^+^ immune cell infiltration

Epithelial damage can often be induced or exacerbated by immune cell infiltrates, and pseudomembranes are often formed by sloughed epithelial cells and intraluminal neutrophils bound in a fibrinous matrix (17, 24). Therefore, immunohistochemical analyses of infected cecal tissues was employed to test GT-dependent effects on MPO^+^ immune cell recruitment, and to determine if epithelial damaged observed in A_GTX_ B_GTX_-infected mice occurred through toxin-host interactions or aggressive immune cell infiltration. The number of MPO^+^ cells per 20x field of view were quantified in single mucosal layers (*n* = 3 animals per treatment). This approach revealed that R20291 and A_GTX_ B+ caused the most significant immune cell influx during disease (∼230 cells per FOV; *p* < 0.001; Fig 5). Although A+ B_GTX_ elicited mild MPO^+^ cell recruitment, it was not significantly higher than that observed in mock-inoculated mice (*p* = 0.2587; Fig 5). Finally, A_GTX_ B_GTX_ caused considerable epithelial damage and pseudomembrane formation, but there were few MPO^+^ immune cells recruited to sites of epithelial damage (Fig 5).

**FIGURE 5.**
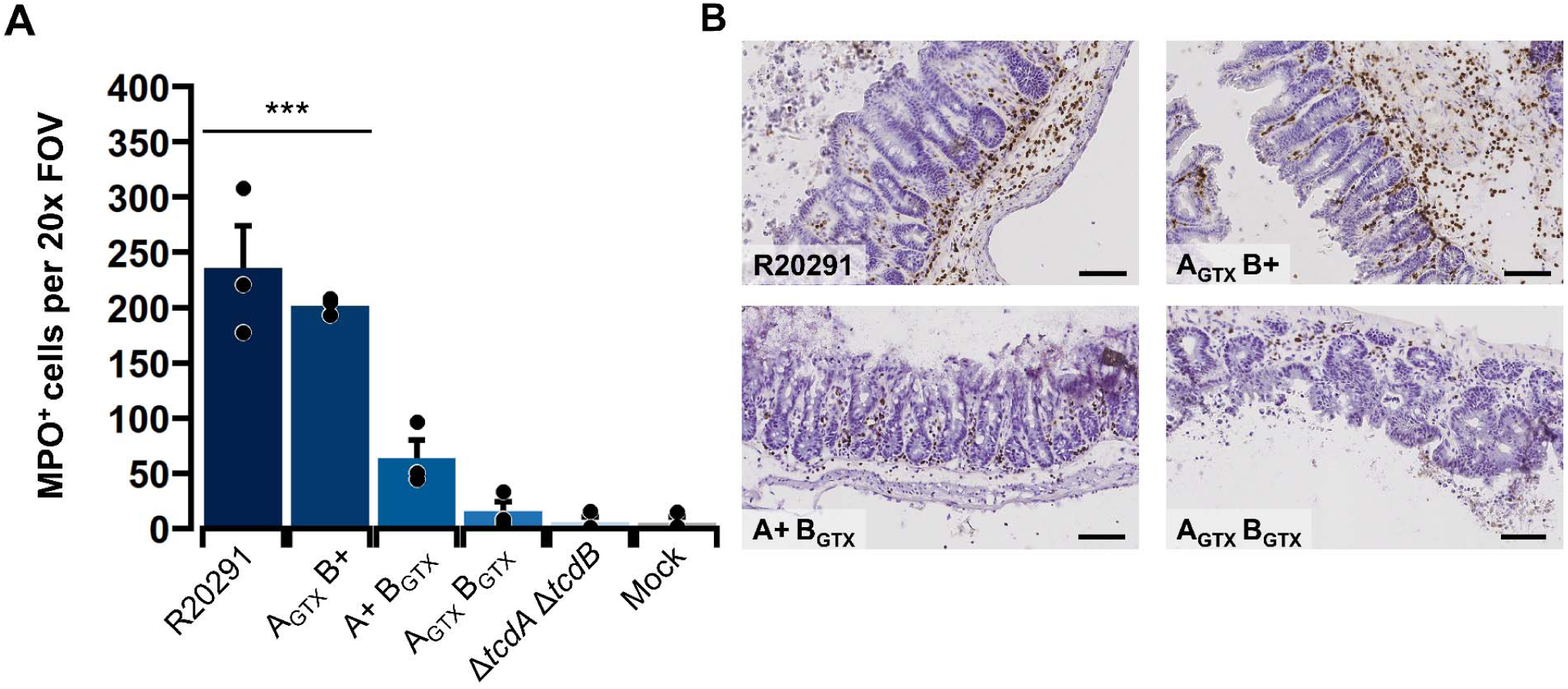
Glucosyltransferase activity of TcdB is required to elicit acute myeloperoxidase-positive immune cell infiltration. (A) Number of MPO^+^ cells per single mucosal layer per 20x field of view (FOV). Each point is the average of three 20x FOVs from one animal. Bars depict the group mean and error bars are the standard error of the mean. Statistical differences between R20291 and A_GTX_ B+ and the other treatments as determined by Tukey’s HSD are shown in a bracket (*\*\*\* p* < 0.001). (B) Representative MPO immunohistochemistry images showing high amounts of immune cell influx into submucosal and mucosal layers in R20291 and A_GTX_ B+, and lower amounts in A+ B_GTX_ and A_GTX_ B_GTX_. Scale bar, 80 μm.

## DISCUSSION

CDI continues to be a significant nosocomial disease in the USA and worldwide, and a more thorough understanding of the function of TcdA and TcdB during infection may open new avenues for prevention and treatment therapies. This includes the development of toxin-inhibiting drugs such as small molecule inhibitors or neutralizing antibodies and nanobodies. These compounds are designed to inhibit specific toxin domains such as the GTD, APD, or the combined oligopeptide repeat (CROPS) domain (5). In recent years, efforts have focused on inhibiting the GT activity of TcdB since TcdB is considered the major driver of severe CDI symptoms. While conceptually an ideal target, multiple investigators have noted that glucosyltransferase-deficient toxin mutants are still cytotoxic or capable of eliciting cytokine responses (9, 12-14, 25, 26).

The role of glucosyltransferase-independent activities during infection are not fully understood. Previous attempts to recapitulate GT-independent effects in murine models have included instillation of recombinant GT-defective TcdB by intrarectal instillation or cecal injection (27, 28). These studies found no GT-independent effects. However, these systems may have lacked the sensitivity of the *C. difficile* infection model, in which toxins are constantly produced and interact with the host epithelium over a course of several days. Recently, GT activity of TcdB was shown to be necessary to cause CDI, and there were no observed GT-independent effects in the mouse model of infection using defined GTX mutants in the *C. difficile* 630 strain (16). This study had two limitations: the first being that the *C. difficile* 630 strain produces few CDI symptoms in the mouse model of infection when compared to epidemic *C. difficile* strains, and it also does not cause epithelial damage (16, 17). The second limitation was that histopathology was assessed at 5 dpi, a timepoint in which mice begin to recover from CDI (16, 17). The aim of our study was to define the GT-dependent and independent effects of *C. difficile* toxins during CDI using the epidemic BI/NAP1/PCR-ribotype 027 R20291 strain in the mouse model of infection and to look for evidence of epithelial damage at the time when weight loss is most severe.

In recent years, TcdB has been accepted as the main virulence factor necessary for fulminant CDI (19, 20). However, TcdA alone can cause mild to moderate CDI symptoms in animal models of infection(19, 20). In this study, mice were inoculated with novel isogenic mutants with deactivated GTDs of either or both toxins, as well as wildtype, double toxin KO, and mock-inoculated controls. The result of these experiments demonstrated that GT activity of TcdA or TcdB alone can cause moderate weight loss during infection, and that GT activity is required for CDI-induced weight loss. Mice inoculated with the wildtype R20291 strain lost the most weight, indicating that the toxins had an additive effect in causing the most severe weight loss outcomes. Additionally, the size of colons and ceca were measured, revealing that GT activity of either toxin was sufficient to cause significant inflammation and subsequent size reduction of both organs. Unexpectedly, we observed that R20291 was the only strain capable of causing severe watery diarrhea, whereas A+ B_GTX_ and A_GTX_ B+ only caused stool to become softer and discolored. Collectively, these results highlight the nuanced and additive effects of each toxin to CDI symptoms including weight loss, diarrhea, and organ inflammation.

Histological damage was assessed in mouse ceca, which is the site of the most severe histopathology in the mouse model of CDI (17). Strains that had functional GT activity of one or both toxins caused severe multifocal edema, whereas GT-inactive or toxin knockout strains caused little to no edema. Inflammation was the most severe in mice inoculated with R20291, A_GTX_ B+, and A+ B_GTX_. However, the severity of inflammation between A+ B_GTX_ and A_GTX_ B_GTX_ was not significantly different, suggesting a mild GT-independent effect on inflammation during infection. Wildtype R20291, A_GTX_ B+, A+ B_GTX_, and A_GTX_ B_GTX_ all caused severe epithelial damage during infection. This phenotype was observed as dead and dying cells sloughing off into the lumen, and by the presence of pseudomembranes. Unexpectedly, Δ*tcdA* B_GTX_ did not elicit epithelial damage different from the double KO and mock controls, which may indicate that GT-independent effects of TcdA are necessary to potentiate epithelial damage during CDI. Indeed, both TcdA and TcdB cause GT-independent effects on immune responses during intoxication *in vitro* and *in vivo* which may synergize to increase the severity of epithelial damage (9, 25, 29). Together, these results demonstrate that edema and inflammation are GT-dependent and are elicited by TcdA or TcdB during infection, although mild inflammation can be caused in a GT-independent manner. They also indicate that epithelial injury occurs through GT-independent mechanisms of TcdA and TcdB during infection.

Since inflammation can contribute to epithelial damage in some models (24, 30–33), immunohistochemical analyses were used to quantify GT-dependent and independent effects on MPO^+^ immune cell influx. Strikingly, high amounts of MPO^+^ cell infiltration were detected in mice inoculated with only R20291 or A_GTX_ B+. In contrast, relatively low levels of MPO^+^ cells were observed in ceca from mice inoculated with A+ B_GTX_ and A_GTX_ B_GTX_ despite their induction of epithelial injury and inflammation. These results indicate that MPO^+^ immune cell infiltration is dependent on GT activity of TcdB, and that A_GTX_ B_GTX_ causes severe epithelial damage and pseudomembrane formation in the absence of MPO^+^ immune cell recruitment.

In conclusion, our data demonstrate that GT activity of either TcdA and TcdB is required for weight loss and organ inflammation, but GT activity of both toxins together causes the most severe weight loss and diarrhea phenotypes. Histological assessment of infected tissues revealed that GT activity of TcdA or TcdB can cause significant edema and inflammation, however GT-activity was not necessary to cause epithelial damage and pseudomembrane formation. Finally, analysis of MPO^+^ immune cell recruitment demonstrated that GT activity of TcdB was required to elicit MPO^+^ immune cells to sites of infection and that GT-independent epithelial damage did not require or elicit MPO^+^ immune cell influx.

While we have observed GT-independent effects during infection, it is possible that therapeutic approaches targeting the GTD will be successful to halt downstream GT-dependent diarrhea and acute inflammatory responses that exacerbate disease severity. However, we must consider that there may be consequences from GT-independent epithelial damage and pseudomembrane formation that might occur when GT activity is neutralized, such as effects on CDI recurrence and/or colon-related diseases. Future studies will aim to elucidate these effects.

## MATERIALS AND METHODS

### *C. difficile* culturing and spore generation

*C. difficile* strains were routinely cultured at 37°C in brain-heart-infusion medium supplemented with 0.5% yeast extract and 0.1% cysteine (BHIS) in an anaerobic chamber (90% nitrogen, 5% hydrogen, 5% carbon dioxide). Strains were stored at -80°C in 20% glycerol for long-term use.

Spores were prepared by transferring a single colony of each strain into 2 mL of BHIS and incubating overnight. The next day, the 2 mL culture was inoculated into 50 mL of Clospore medium, which was then grown for 10 days anaerobically (34). After sporulation, the suspension was centrifuged at 4,000 x g at 4°C and washed five times in cold sterile water. Spores were suspended in 1 mL sterile water and heat treated at 65°C for 20 minutes to eliminate vegetative cells. Viable spores were quantified through serial dilutions and spotting on BHIS + 0.1% taurocholate (TA) plates. Spore stocks were stored at 4°C until use.

### *C. difficile* mutant generation

To study GT-dependent and independent effects during infection, *C. difficile* strains with mutations deactivating the GT catalytic domain (“GTX”) of TcdA (TcdA::D285N/D287N), TcdB (TcdB::D286N/D288N), or both toxins (TcdA::D285N/D287N-TcdB::D286N/D288N) were generated (Table 1). Mutations were introduced in *tcdA* and *tcdB* genes of the BI/NAP1/PCR-ribotype 027 epidemic *C. difficile* R20291 strain using homologous allelic exchange (22). Four novel strains were generated for this study: A_GTX_ B+, A+ B_GTX_, A_GTX_ B_GTX_, and Δ*tcdA* B_GTX_ (Table 1).

Mutants with deactivated catalytic domains were created by PCR amplifying a 2393 bp fragment of *tcdA*, and a 2332 bp fragment of *tcdB* from *C. difficile* R20291 genomic DNA (gDNA), then cloning amplicons into separate pMTL-YN4 vectors at flanking *AscI* and *SbfI* restriction sites (22). Single nucleotide mutations (SNMs) in *tcdB* were generated using Q5 Site-Directed Mutagenesis according to manufacturer’s protocol (New England Biolabs). Briefly, oligonucleotides that contained the SNMs were used to PCR amplify the entire pMTL-YN4::*tcdB* vector (7788 bp), then treated with Kinase-Ligase-*DpnI* (New England Biolabs) prior to heat shock transformation into chemically competent DH5α *Escherichia coli* cells (Table 1, 2). Q5 Site-Directed Mutagenesis did not work for *tcdA*; therefore, Gibson assembly was used as an alternative approach. Each half of the pMTL-YN4::*tcdA* vector (7816 bp total) was amplified in two separate reactions using oligonucleotides containing SNMs in the overlap region to generate the SNMs in *tcdA* (Table 2). Finally, to generate the *tcdA* knockout (KO) cassette, *C. difficile* R20291 gDNA was PCR amplified with oligonucleotides flanking the *tcdA* gene that include the start and stop codons (Table 2). To create a whole-gene deletion, left and right homology arms (1039 bp and 1017 bp, respectively) of the amplified *tcdA* were created and joined together by Gibson assembly into the pMTL-YN4 plasmid at *AscI* and *SbfI* restriction sites to create pMTL-YN4::Δ*tcdA*, which was introduced into DH5α *E. coli* via heat shock (Table 1, 2) (22).

**TABLE 2.**
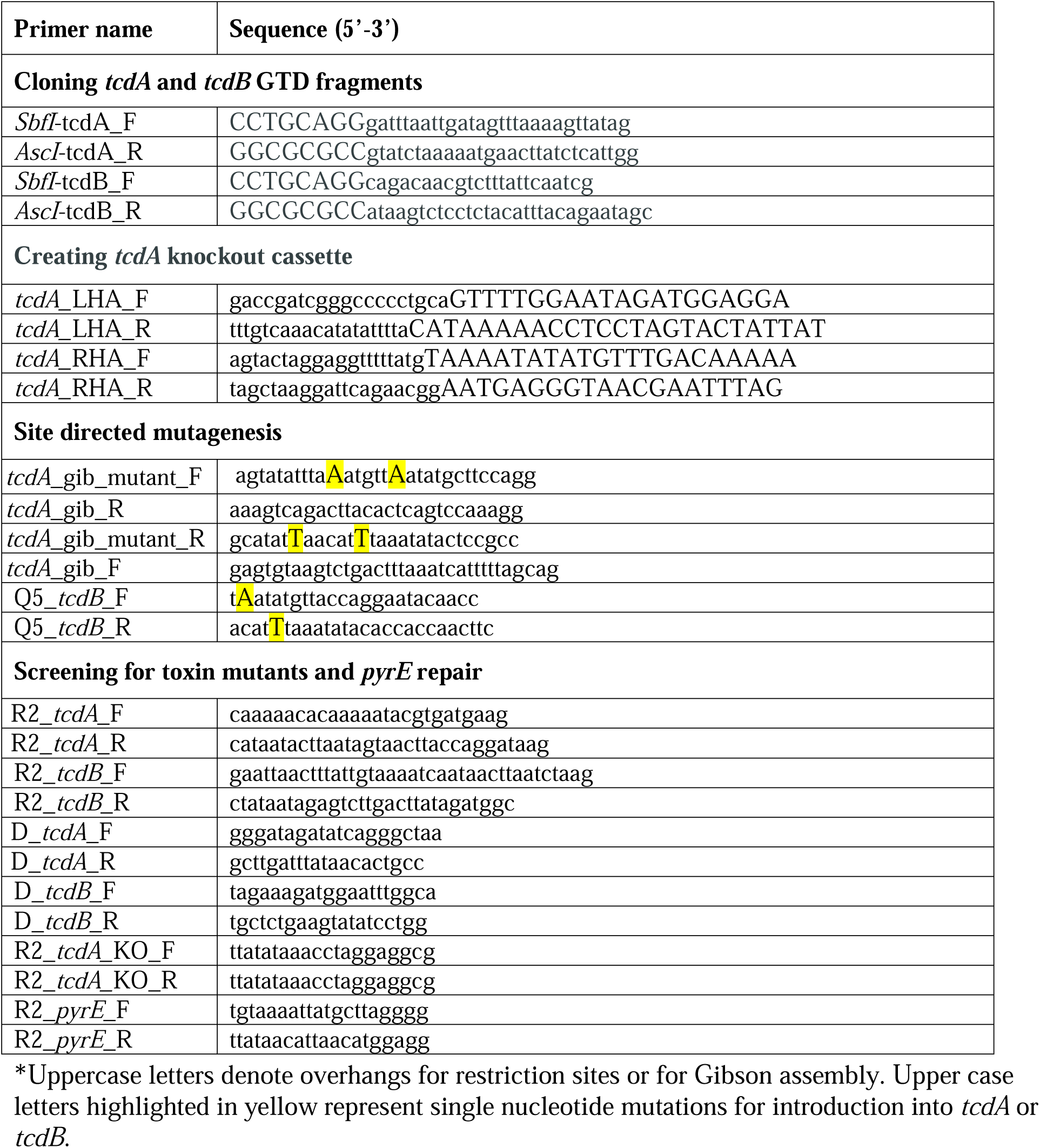
Oligonucleotides used in this study.

*E. coli* containing pMTL-YN4::*tcdA*, pMTL-YN4::*tcdB*, or pMTL-YN4::Δ*tcdA* were conjugated into a *C. difficile* R20291Δ*pyrE* mutant strain as previously described (Table 1) (22, 35). Transconjugants were selected based on thiamphenicol (15 μg/mL) resistance on BHIS plates supplemented with D-cycloserine (250 μg/mL) and cefoxitin (8 μg/mL) (22). Single crossover integrants that were thiamphenicol resistant were further identified by PCR amplification and Sanger sequencing. Confirmed single cross-over clones were grown on BHIS supplemented with 500 μg/mL 5-fluoroorotic acid and 1 μg/mL uracil to induce *pyrE*-based counter selection for double crossover mutants. Plasmid loss was then confirmed by thiamphenicol sensitivity. Single and double toxin mutants, as well as *tcdA* KO was confirmed by PCR amplification and Sanger sequencing of each site (Table 2). After sequence confirmation, the *pyrE* locus of each strain was restored to wildtype as previously described (22).

### Whole genome sequencing and mutant confirmation

Whole genome sequencing was performed to confirm the presence and absence of mutations for all strains used in this study (Table 1). Strains were cultured overnight in 3 mL liquid BHIS, then gDNA was extracted using the MasterPure Gram Positive DNA Purification kit according to manufacturer’s protocol (Lucigen). Library preparation and sequencing was performed by the Microbial Genome Sequencing Center (MiGS). Briefly, DNA quality and quantity was assessed with a Qubit fluorometer prior to library preparation using the Illumina DNA Library Prep kit according to manufacturer’s protocol. Samples were sequenced on the Illumina NextSeq 2000 platform as paired end 2×150 bp reads to generate 200 Mbp total reads.

Quality control of reads and adapter trimming was performed using instrument software. Each genome underwent *de novo* assembly and annotation using SPAdes and RASTtk, respectively, in PATRIC 3.6.9 (36–38). Genomes of all strains were aligned to the reference *C. difficile* R20291 genome (NC_013316) (39) using Burrows-Wheeler Aligner (BWA-mem), then single nucleotide variants and indels were analyzed using FreeBayes in PATRIC 3.6.9 (38, 40, 41). Finally, *tcdA* and *tcdB* genes from all strains were aligned and presented using Clustal Omega (42). Sequencing reads were deposited to NCBI Sequencing Read Archive accession number: PRJNA762329.

### *C. difficile* toxin production and secretion *in vitro*

Immunoblotting of *C. difficile* cell pellets and culture supernatants was used to test toxin production and secretion in the wildtype and mutant strains. All strains were grown overnight on BHIS + 0.1% TA plates from glycerol stocks. The next day, individual colony forming units (CFUs) from each strain were transferred into 3 mL BHIS and cultured for 18 hours. Optical density at 600 nm (OD_600_) was then measured, and each culture was diluted to OD_600_ = 0.01 in 3 mL of Tryptone Yeast (TY) medium and grown for 24 hours. OD_600_ was again measured, and cultures were adjusted to OD_600_ = 1 in 1 mL of TY. Bacterial suspensions were centrifuged at 13,000 x g for 15 minutes at 4°C, then the supernatants were filtered through a 0.2 μm cellulose acetate membrane (VWR) into sterile 1.5 mL Eppendorf tubes. Bacterial pellets were resuspended in 100 μl PBS pH 7.4, and all pellets and supernatants were flash frozen in liquid N_2_ until use.

Tubes containing bacterial pellets and supernatants were thawed on ice, and 20 μl of pellet and supernatant suspensions were transferred to tubes containing 10 μl Laemmli buffer and β-mercaptoethanol then boiled at 90°C for 10 min. Five microliters of 6x loading dye were added to each tube, and equal volumes of each suspension were loaded into two sodium dodecyl sulphate-polyacrylamide gels (SDS-PAGE; 4-20% Mini-PROTEAN TGX Protein Gels, Bio-Rad) and run at 150 V for one hour. One gel was used for Western blotting and the other was stained with SYPRO Ruby gel stain (Bio-Rad) for visualizing total protein.

Proteins were transferred from the SDS-PAGE gel to a polyvinylidene fluoride membrane at 100 V for 1 hour. The membrane was then transferred to a container and blocked in 5% milk in tris buffered saline pH 7.6 supplemented with 0.1% Tween 20 (TBST) for 30 minutes at 4°C. After blocking, the solution was removed, and the membrane was incubated in TBST-5% milk containing the anti-TcdB-GTD antibody, 20B3, at a 1:1000 dilution overnight at 4°C (12, 43). The next day, the membrane was washed three times in TBST then incubated with goat-anti-mouse Horse Radish Peroxidase (HRP) diluted 1:10,000 in TBST-5% milk solution for 1 hour at 4°C. The membrane was washed three more times in TBST before addition of Immobilon Western Chemiluminescence HRP Substrate (Millipore) and visualization via film development. After imaging, the membrane was stripped in a harsh SDS and β-mercaptoethanol-based buffer for 30 min. The membrane was re-blocked and probed with a 1:1000 diluted anti-TcdA antibody (clone PCG4.1, Novus Biologicals), then processed using the exact same method described for TcdB.

### *C. difficile* growth *in vitro*

Strain growth in artificial media was measured to test differences in growth rate *in vitro*. Briefly, each strain was plated on BHIS from glycerol stocks, then grown overnight as described above. Five CFUs from each strain were transferred into 3 mL BHIS and grown statically for 18 hours. The next day, cultures were diluted to OD_600_ = 0.05 in wells of a 96-well plate (Nunc) containing 200 μl BHIS. Control mock-inoculated wells were included by adding sterile PBS into six wells. OD_600_ was measured every hour for 24 hours in a BioTek Synergy 4 microplate reader (BioTek-Agilent). *In vitro* growth graphs were generated using ggplot2 and ggprism, and statistical differences between groups at each timepoint were assessed by calculating and plotting the 95% confidence interval in R v. 4.0.3.

### Mouse model of *C. difficile* infection

This study was approved by the Institutional Animal Care and Use Committee at Vanderbilt University Medical Center (VUMC) and performed using protocol M1700185-01. Our laboratory animal facility is AAALAC-accredited and adheres to guidelines described in the Guide for the Care and Use of Laboratory Animals. The health of the mice was monitored daily, and severely moribund animals were humanely euthanized by CO_2_ inhalation followed by cervical dislocation. All animals in this study were C57BL/6J female mice between 8-10 weeks of age purchased from Jackson Laboratories. Mice were assimilated to the new facility for one week prior to antibiotic treatment to reduce stress. Mice were housed in a pathogen-free room with 12-hour cycles of light and dark. Cages were changed every two weeks to ensure clean bedding, and they had free access to food and water.

Differences in virulence between mutant and wildtype strains were tested using the cefoperazone mouse model of *C. difficile* infection (17). Nine-week-old C57BL/6J female mice were treated with cefoperazone (0.5 mg/kg) in sterile drinking water *ad libitum* for five days, followed by two days of untreated water prior to inoculation (17). Mice were inoculated with 10^5^ CFU/mL spores in 100 μl of sterile PBS via transoral gastric gavage.

Mouse weight and symptoms were recorded daily, and daily stool samples were collected. Stool samples were scored on a 1-4 scale for color and composition as an additional metric of disease, where 1 = normal, well-formed stool; 2 = well-formed, discolored stool; 3 = moist, soft, and discolored stool; and 4 = wet tail, watery diarrhea, and empty cecum & colon. Stool was then weighed, macerated in 500 μl sterile PBS pH 7.4, and 5-fold dilution plated on TA (10% w/v), D-cycloserine (10 mg/mL), cefoxitin (10 mg/mL), and fructose agar (TCCFA) semi-selective media to enumerate *C. difficile* titers *in vivo*. Weight loss, stool score, and daily CFU g^-1^ graphs were plotted with ggplot2 and ggprism, and statistical differences at each timepoint were tested using one-way ANOVA and Tukey’s HSD test (weight loss) or Kruskal-Wallis and Dunn’s test (CFU g^-1^ stool) in R v. 4.0.3.

Mice were humanely euthanized at two- or four-days post-inoculation by CO_2_ inhalation. Post-sacrifice, the colon and cecum were excised from each animal and imaged to measure size. After imaging, the whole colon was flushed with sterile PBS, cut transversally, then splayed open and fixed in 2% paraformaldehyde (PFA) at 4°C for 2 hours. Intact ceca were laid flat on sterile Whatman filter paper and fixed in 2% PFA for two hours at 4°C. After fixation, tissues were washed three times in cold, sterile PBS, then transferred to an ice-cold solution of 30% sucrose, 1% sodium azide and incubated for 16 hours at 4°C. Tissues were then embedded in OCT (Optimal Cutting Temperature embedding medium; Fisher Healthcare) on dry ice-cooled ethanol and stored at -80°C until use. Images were used to determine the length of colons and area of ceca (*n* = 6 per treatment) using ImageJ. These metrics were graphed using ggplot2 and ggprism, and statistical differences were determined using one-way ANOVA and Tukey’s HSD test in R v. 4.0.3.

### Histopathological tissue assessment

Frozen tissue blocks of ceca and colons were sectioned on a Leica CM1950 Cryostat (Leica Biosystems) as 7 μm slices onto Superfrost Plus microscope slides (Fisher Scientific) and stored at -80°C until use. To assess histopathology, cecum sections were stained with hematoxylin & eosin (H&E; Vector Labs), and conditions were masked for a board-certified gastrointestinal pathologist to score edema, inflammation, and epithelial damage based on published criteria (*n* = 6-9 per treatment) (17). Histological scores were graphed with ggplot2 and ggprism, and statistical differences were determined using one-way ANOVA and Tukey’s HSD test in R v. 4.0.3. Presented images were captured using a Leica SCN400 Slide Scanner automated digital image system from Leica Microsystems at the Digital Histology Shared Resource (DHSR) at VUMC. Whole slides were imaged at 40x magnification to a resolution of 0.25 μm/pixel.

### Assessment of myeloperoxidase-positive immune cell recruitment during infection

Immune cell infiltration was detected by anti-myeloperoxidase (MPO) staining performed at the VUMC Translational Pathology Shared Resource. Briefly, slides were placed on the Leica Bond RX IHC stainer. Slides were placed in a Protein Block (DAKO) for 10 minutes, then incubated with anti-MPO (DAKO A0398) for one hour at 1:4000 dilution. The Bond Polymer Refine detection system was used for visualization. Slides were then dehydrated, cleared and coverslipped. To quantify immune cell infiltrates, a single mucosal layer was visualized in a 20x field of view (FOV) at a time, and the number of MPO^+^ cells were counted per FOV in that region. The number of MPO^+^ cells was quantified in three FOVs per mouse per treatment. The average MPO^+^ cells per mouse per treatment (*n* = 3) were plotted using ggplot2 and ggprism, and differences were tested using one-way ANOVA and Tukey’s HSD test in R v. 4.0.3. Presented images were captured using a Leica SCN400 Slide Scanner at the VUMC DHSR as described above.

## ACKNOWLEDGEMENTS

This work was supported by NIH AI957555, VA BX002943, and T32 DK007673. Thank you to Melissa Farrow and members of the Lacy lab for stimulating discussions and feedback on these experiments, to Nick Markham and members of the Skaar lab for advice on the mouse model of *C. difficile* infection, and to Grace Morales for advice on genome analyses. Also, thank you to Nigel Minton for supporting this work by providing lab space and equipment access for Rory Cave.

---

**SUPPLEMENTAL FIGURE 1.**
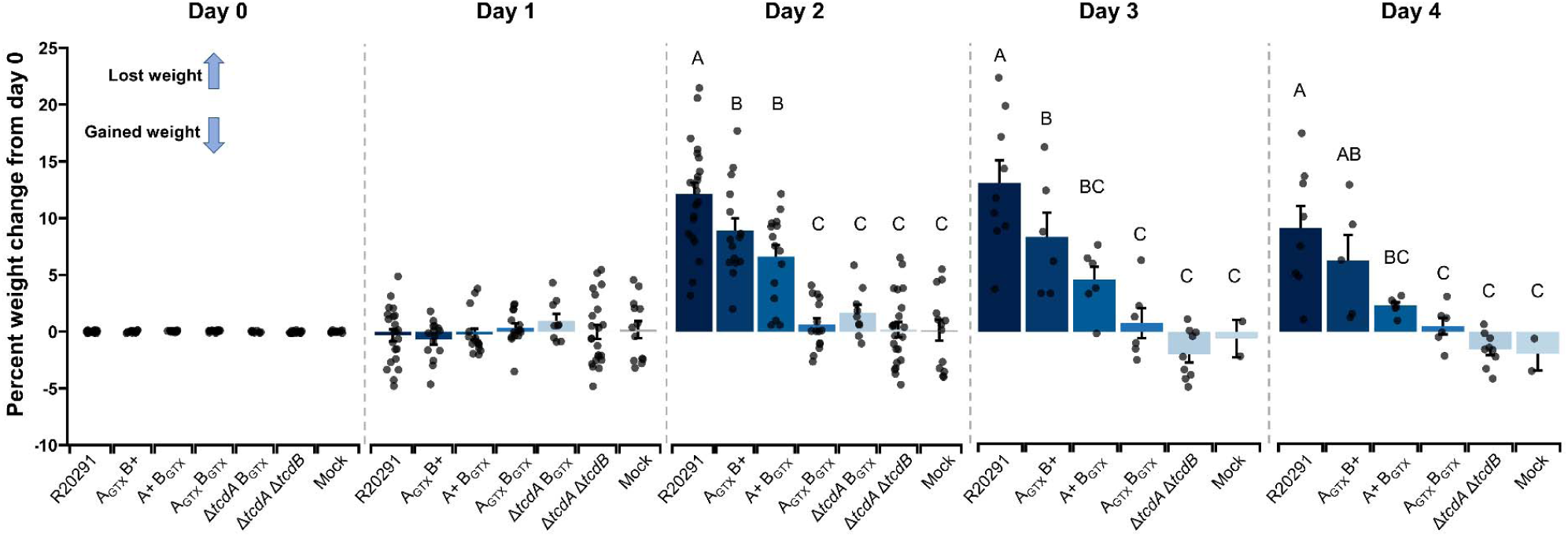
Percent weight loss from day 0 weight during infection studies. Positive increases signify weight lost and negative decreases indicate weight gained. Each point is an individual animal and bars represent group means at each day. Error bars denote standard error of the mean. Significantly different groups as determined by Tukey’s HSD test are shown using letters where groups containing multiple letters are not significantly different to individuals containing those single letters (*p* < 0.05).

## Notes

### Competing Interest Statement

The authors have declared no competing interest.

